# Variability in cadmium accumulation in quinoa grain: potential role of crop domestication and other factors mediating differences in resilience and uptake

**DOI:** 10.1101/2025.06.30.662325

**Authors:** Alejandro Rodriguez-Sanchez, Martha Jimenez-Castaneda, Timothy Filley, David Brenner, Dennis Macedo, Victor H. Casa, Lori Hoagland

## Abstract

Quinoa (*Chenopodium quinoa*) is a nutritious crop expanding in importance worldwide, though its capacity to accumulate cadmium (Cd) can put consumers at risk. The objective of this study was to identify genotypes that vary in Cd uptake and explore how domestication, saponins, and root-associated microbes could influence these differences. Twelve genotypes representing wild ancestors and modern landraces or cultivars from four distinct ecoregions were grown in a greenhouse trial using potting media amended with Cd. Differences in phenological developmental traits were recorded during growth, and plant biomass, microbial derived amino sugars in roots, and concentrations of Cd and other essential elements in grains were quantified at harvest. Phenological development of Ancestors were more adversely affected by Cd than modern genotypes, and Cd concentrations were higher in those developed later near coastal regions in South America and the United States, suggesting that quinoa become more tolerant over domestication and selection. Concentrations of several elements were also lower in the Ancestors vs. modern genotypes in response to Cd, indicating that modern genotypes may not downregulate root transporters as an adaptation to Cd stress. Fungal derived amino sugars were negatively correlated with Cd concentrations in the Coastal ecotypes which had the highest uptake, suggesting that the presence of root-associated fungi could help restrict translocation aboveground. Results of this study will provide breeders with new insights into factors that could aid in selecting for quinoa with low Cd-uptake. Early genotypes selected in the Altiplano region may be most useful in these breeding programs.

## 1. Introduction

The pseudo-cereal quinoa (*Chenopodium quinoa*) has become an important crop marketed worldwide due to its high-quality nutrition at a relatively low cost (FAO, 2011). Recognizing its significance, the Food and Agriculture Organization (FAO) declared 2013 the “International Year of Quinoa” (FAO, 2013). Quinoa seeds or grains contain high levels of protein, lipids, fibers, vitamins, minerals and amino acids including lysine, among other beneficial compounds (de Lima Brito et al., 2022). Consequently, the grain is included in a wide variety of products including many baby foods due to its nutritional profile and fact that it is also gluten-free (Ayseli et al., 2020; Jimenez et al., 2020; Venlet et al., 2021). Unfortunately, quinoa plants can bioaccumulate several toxic heavy metals and metalloids such as arsenic (As), lead (Pb), and cadmium (Cd) in its edible grains (Pizarro et al., 2016; Haseeb et al., 2018; Shabbir et al., 2020; Gardener et al., 2018; Gardener et al., 2019). The presence of elevated concentrations of these contaminants can pose a threat to human health (Roman-Ochoa et al., 2021), especially among infants and young children who are at higher risk (Gardener et al., 2018; Gardener et al., 2019; ECP Baby Food Staff Report, 2021; Flannery et al., 2022; Amadi et al., 2022). Ingestion of high amounts of Cd is particularly problematic, since this heavy metal can lead to osteomalacia, cardiovascular diseases, kidney damage, and bladder and prostate cancer, among other health disorders (Witkowska et al., 2021). Cd also tends to be highly mobile in soil (Shahid et al., 2017), making it difficult to prevent plant uptake even when using soil amendments like biochar that can bind this element and reduce its bioavailability in soil (Zea et al., 2022). Therefore, development of new strategies for reducing Cd in quinoa grains is critical to ensuring the continued success of this important crop.

One way to help reduce human health risks is to develop quinoa cultivars that can restrict Cd uptake. Differences in Cd bioaccumulation among quinoa genotypes has previously been demonstrated (Bhargava et al., 2008), and further efforts to identify the mechanisms mediating these differences have potential to aid in the development of new commercially relevant varieties that can reduce human health risks. For example, plants can limit Cd uptake by shutting down root transporters (Sun et al., 2021), though how this can influence the uptake of other essential plant elements like iron (Fe) and zinc (Zn), which are important components of quinoa’s nutritional quality, are still unclear. Other factors that could influence differences in Cd bioaccumulation include the presence of saponin compounds in grains since these compounds can bind to heavy metals (Nobahar et al., 2021), and are often present in high concentrations within quinoa pericarps and to a lesser extent in seed coats (Gonzalez, 1996, Roman-Ochoa et al., 2023). This is tied to *Chenopodium* seed anatomy, in which wild accessions usually have thicker seed coats than domesticated accessions (Sukhorukov & Zhang, 2013), with thickness correlated with saponin content (Hazzam et al., 2020). Interaction with root-associated microbes that can alter Cd bioavailability in soil and/or sequester Cd in structures such as variable N-rich biopolymers within their membranes could also reduce Cd uptake (Luo et al., 2019).

Understanding how these plant traits evolved over domestication can provide important clues and resources to help plant breeders select for improved cultivars. For example, maize (*Zea mays*) and durum wheat (*Triticum duram*) have been shown to accumulate higher levels of heavy metals as a result of domestication (Vielle-Calzada et al., 2010; Macaferri et al., 2019), indicating that wild crop ancestors or land races could provide valuable sources of germplasm for developing cultivars with lower heavy metal uptake. Crop domestication gradients have also been shown to influence plant root interaction with beneficial soil microbes (Jaiswal et al., 2020), as well as root transporter activity (Cao et al., 2018), and consequently could be linked to accumulation of Cd in the grain.

Modern quinoa plants are believed to have evolved from a wild tetraploid of *Chenopodium berlandieri* var. *zschakei*, native to North America (Bazille et al., 2013). Seeds from this species are thought to have been transported to the Andes mountains via bird dispersal or human migration. In the Andes, the plant underwent further evolutionary changes, leading to the development of *Chenopodium berlandieri* var. *berlandieri* and *Chenopodium hircinum*, which were eventually domesticated into *C. quinoa* (Bazille et al., 2013; Jarvis et al., 2017).

In South America, it is believed that the first domesticated quinoa plants originated in highland or Altiplano regions near Lake Titicaca between Peru and Bolivia. From there, the plants were grown in the Pacific coastal areas of Peru and Chile in successive domestication events (Jarvis et al., 2017). At the same time, *Chenopodium berlandieri* var. *zschakei* is thought to have evolved into *Chenopodium berlandieri* var. *sinuatum* in southern Mexico, which subsequently gave rise to *Chenopodium berlandieri* subs. *nuttalliae* (Bazille et al., 2013).

The objective of this study was to identify quinoa genotypes that vary in Cd resilience and uptake and investigate potential factors that could influence these differences. Specifically, we aimed to 1) quantify how domestication could influence differences in how quinoa plants tolerate Cd stress via changes in phenological development and elemental uptake; 2) determine whether differences in the amount of saponins present in quinoa seeds could influence Cd bioaccumulation in grains; and, 3) investigate how interaction with root-associated microbes that can regulate elemental cycling and uptake could influence these differences.

## 2. Materials and methods

### 2.1 Selection of Chenopodium accessions

Using the current paradigm for the domestication of quinoa, we selected twelve (12) *Chenopodium* genotypes representing the domestication process including wild quinoa relatives, intermediate relatives or landraces, and modern cultivars developed in unique ecoregions of South America and the U.S. (Figure 1; Table 1). Some of the genotypes were also selected based on previous reports of differences in Cd uptake (Bargava et al., 2008) and saponin concentration in seeds (Medina-Meza et al., 2008). All seeds were acquired from the USDA-ARS North Central Regional Plant Introduction Station (NCRPIS) based in Ames, Iowa, which serves as the U.S. germplasm repository for *Chenopodium*.

**Figure 1.**
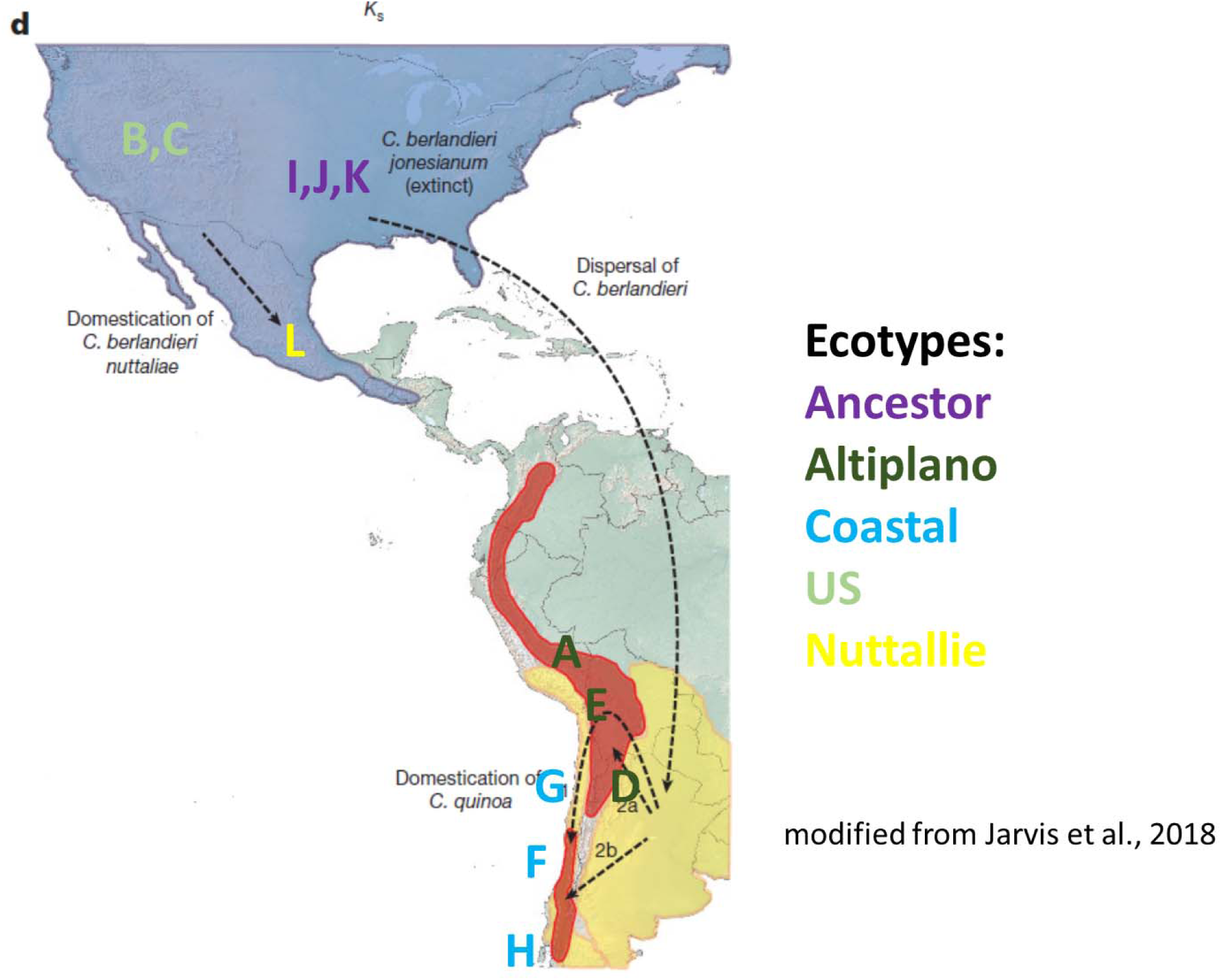
Geographical distribution of *Chenopodium* genotypes included in this study. Genotypes I, J and K are the Ancestor genotypes; genotypes A, E and D are altiplano; genotypes F, G and H are coastal; and B and C come from altiplano varieties bred in the United States.

**Table 1.**
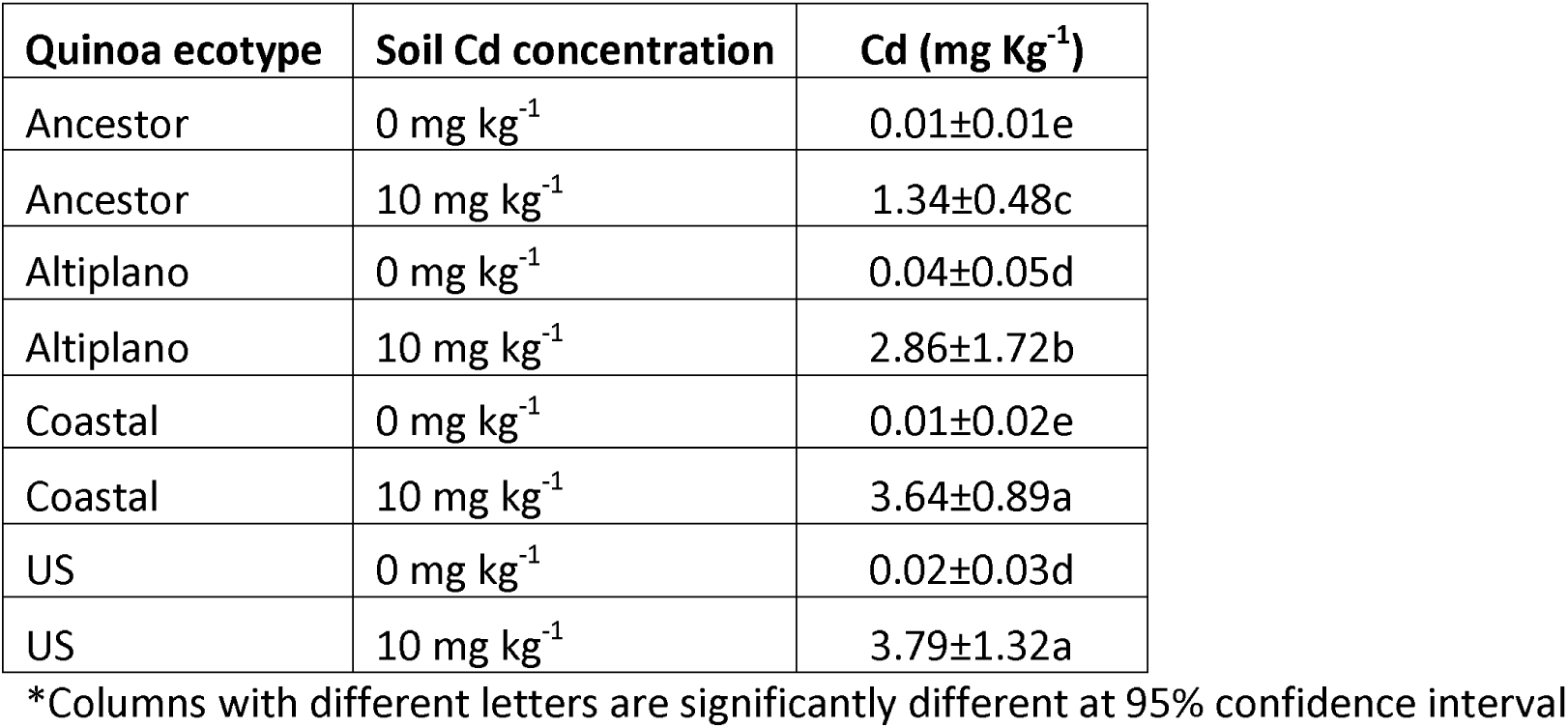
Cd concentration in seeds of the different quinoa ecotypes

| Quinoa ecotype | Soil Cd concentration | Cd (mg Kg<sup>-1</sup>) |
| --- | --- | --- |
| Ancestor | 0 mg kg<sup>-1</sup> | 0.01±0.01e |
| Ancestor | 10 mg kg<sup>-1</sup> | 1.34±0.48c |
| Altiplano | 0 mg kg<sup>-1</sup> | 0.04±0.05d |
| Altiplano | 10 mg kg<sup>-1</sup> | 2.86±1.72b |
| Coastal | 0 mg kg<sup>-1</sup> | 0.01±0.02e |
| Coastal | 10 mg kg<sup>-1</sup> | 3.64±0.89a |
| US | 0 mg kg<sup>-1</sup> | 0.02±0.03d |
| US | 10 mg kg<sup>-1</sup> | 3.79±1.32a |

\*Columns with different letters are significantly different at 95% confidence interval

### 2.2 Greenhouse experiment

Greenhouse experiments were conducted at Purdue’s Horticulture Plant Growth Facility in West Lafayette, IN, using twelve *Chenopodium* genotypes (Table 1). Plants were grown in one-gallon pots filled with BM7 potting media (www.berger.ca). Cd was added to the potting media at three concentrations (0, 5, and 10 mg/kg) using a cadmium chloride (CdCl_2_) solution, with six replicates per genotype for each Cd concentration. The potting media was conditioned for 30 days to allow Cd to bind to the soil particles before planting. During this incubation period, the pots were irrigated with tap water to achieve 50% moisture holding capacity and help distribute the Cd within the media profile but prevent leaching. After the incubation period, five seeds of each *Chenopodium* genotype were planted in each pot and the pots were thinned to one seedling per pot following germination.

The pots were arranged in a randomized complete block design within the greenhouse and subject to the following greenhouse conditions: photoperiod of 14 h, average day-time and night-time temperatures of 22 °C and 18 °C ± 1 °C, respectively, and relative humidity maintained between 50-70%. All pots were fitted with drippers and irrigated as necessary with water supplemented with water-soluble fertilizer (20N-1.3P-15.8K; ICL Specialty Fertilizers, Dublin, OH) to provide the following (in mg/L): 150 N, 9.8 P, 119 K, 12 Mg, 21 S, 1.5 Fe, 0.4 Mn and Zn, 0.2 Cu and B, and 0.1 Mo. Nitrate and ammoniacal sources of nitrogen (N) were provided as 61% and 39% total N respectively. The experiment was terminated after all plants had mature seeds, which occurred approximately 134 days after seeding. The genotype L (*C. berlandieri* subsp. *nuttalliae*), did not produce any seeds under the conditions in our plant growth facility, and genotype D (*C. quinoa*; accession #PI587173) produced insufficient seeds, thus, these genotypes were excluded from subsequent assays for quantifying elemental uptake or amino sugars described below.

### 2.3 Chenopodium phenological growth stages

The plants were monitored on a daily basis and the time to reach key phenological growth stages (inflorescence emergence, flowering, complete pollination or fruit development, and ripening) were recorded following the Biologische Bundesanstalt, Bundessortenamt und Chemische Industrie (BBCH) scale developed for quinoa (Sosa-Zuñiga et al., 2017). Plant height was measured 28 and 56 days after seeding using a ruler. Chlorophyll content was estimated at three time points (28 or 56 days after seeding and harvest) using a SPAD-502 meter (Konica Minolta, Inc., New Jersey, USA).

### 2.4 Chenopodium harvesting and seed processing

At harvest, root and shoot length were measured using a ruler, and the fresh shoot and root weight were recorded before and after oven-drying at 60 °C. Seeds were collected at maturity and stored at room temperature until processing to quantify yield and prepare seeds for elemental analyses. Impurities present with the *Chenopodium* seeds (e.g. chaff) were removed using seed pan cleaners (minimum size of 1/16 inches). For *C. berlandieri* genotypes characterized by small seed size, a Carter-Day fractioning aspirator (Carter-Day, US) was used to clean the grains. All seeds were then ground using a UDY grinder Seedburo 3010SM/C (Seedburo, US) with 0.25 mm grid. The grinding of the seeds did not discern pericarp from seed coat.

### 2.5 Determination of macro- and micronutrients and Cd in the seeds

Seed powders from plants grown at 0 and 10 mg/Kg Cd concentration were digested in duplicate (0.5 g each) reactions using 70% HNO_3_ (10 mL) in a MARS 6 microwave digester and Xpress vessels (CEM Corporation, Charlotte, NC) at 200 °C, 800 psi, and a power of 900-1050 watts.

After diluting the digests to 2% HNO_3_, total concentrations of Cd as well as boron (B), calcium (Ca), cobalt (Co), copper (Cu), iron (Fe), magnesium (Mg), manganese (Mn), nickel (Ni), phosphorous (P), potassium (K), sodium (Na), silicon (Si), sulfur (S) and zinc (Zn) in the seed powders were determined on Purdue University’s campus using an Inductively Coupled Plasma- Optimal Emissions Spectroscopy (ICP-OES) (Shimadzu ICPE-9820, Kyoto, Japan) with a nebulizer to increase accuracy. The periodic table mix 1 for ICP, TraceCERT grade for 33 elements including Cd (Sigma Aldrich, Burlington, MA, US) was used as a standard reference. Blanks and additional standard checks were run periodically as a quality assurance measure.

Additional information regarding quality control measures to quantify elemental concentrations in plant tissues can be found in Table S1. *Chenopodium* seeds grown in 10 mg/Kg Cd were also sent to Brooks Applied Labs (Seattle, WA) to verify Cd concentrations using Inductively Coupled Plasma-Mass Spectroscopy (ICP-MS) using their standard protocols (EPA 1600-series for low level metals analysis). Brooks Applied Laboratory is accredited for quantification of trace amounts of heavy metals in food matrices (https://brooksapplied.com/resources/certificates-permits/).

### 2.6 Chemical analysis of fine roots

Dry secondary roots were powdered using a SPEX 6750 cryogenic grinder (Spex Centriprep Inc., US). Because of low volumes, the powders from individual *Chenopodium* genotypes were carefully mixed and combined into one of four ecotypes (Table 1). The powder obtained from homogeneous samples was used for subsequent analyses.

The carbon (OC) content, nitrogen (N) content, δ^13^C, and δ^15^N values of the root samples were determined in duplicate using a Sercon Ltd (Crewe, UK) EA-CN1 elemental analyzer interfaced to a Sercon 20/22 isotope ratio mass spectrometer (IRMS, Crewe, UK). The δ^13^C and δ^15^N values are expressed relative to the international Vienna Pee Dee Belemnite (VPDB) and air standard, respectively (Coplen, 2011). Analytical precision of δ^13^C and δ^15^N values of the laboratory working standards, NIST 1547, USGS 61, Acetanilide and maize were <0.2‰ and <0.22‰, respectively. The carbon to nitrogen (C:N) ratio of the roots was determined by dividing the %C content by the %N content for each ecotype.

Microbial-derived amino sugars present in the roots, including galactosamine (GalN), glucosamine (GlcN), muramic acid (MurA), and mannosamine (ManN) were extracted in duplicate, using 20 µg of the root sample following a modification of the protocol established by Zhang and Amelung (1996) and Liang et al. (2012). Analyses were carried out using a Shimadzu 2010 gas chromatography system equipped with an AOC-20i Shimadzu auto-sampler and programmable temperature vaporization (PVT) inlet, interfaced to a Shimadzu GCMS- QP2010 Plus mass spectrometer operated in electron ionization mode using He as the carrier gas at a constant flow (see Appendix 1 for a detailed description and calculations).

### 2.9 Statistical analyses

Differences in Cd concentrations in the grain of plants grown in 10 mg kg^-1^ potting media and quantified using ICP-MS were evaluated with SAS 9.4 (SAS Institute, Cary NC) statistical software using a one-way analysis of variance (ANOVA); subsequent multiple comparisons (t-tests) were conducted using the Tukey’s honest significance test.

To evaluate differences among the other growth parameters (i.e. SPAD, length and weight of roots and shoots at harvest, root:shoot ratios, phenological stages, and concentrations of Cd and other elements quantified using ICP-OES), the results from individual genotypes were combined into one of four ecotypes (Table 1) to be consistent with the chemical analyses of *Chenopodium* roots. All statistical analyses for these parameters were computed using ANOVA-based pairwise Mann-Whitney U tests using PAST v3.1 software (Hammer et al., 2006). Finally, correlations between each of these parameters and Cd concentrations present in the seeds at harvest were evaluated using Pearson’s ρ correlation computed by PAST v3.1 software.

To test the effect of Cd-rates on the concentration of root- carbon (OC), root nitrogen (N), and amino sugar derived C and N, one-way ANOVA analyses were performed in combination with a post hoc separation of means by Fisher’s least significant differences (LSD). RStudio software (version 1.4.1106) was used for these statistical procedures.

## 3. Results

### 3.1 Effect of Cd concentration and ecotype on phenological development

PERMANOVA results suggest a differentiating effect caused by ecotype (*F*=24.191, *p*=0.0001) and Cd concentration (*F*=3.0427, *p*=0.0195), but not for their interaction (*F*=0.68468, *p*=0.172). When comparing relationships between Cd concentration and individual development parameters and growth stages, correlations varied among the four *Chenopodium* ecotypes (Figure 2). Specifically, the Ancestor ecotype had lower SPAD readings 28 days after seeding and lower plant height at 28 and 56 days after seeding in response to Cd. In contrast, only plant height at 56 days after seeding was influenced by Cd in the Altiplano ecotype, and inflorescence emergence occurred earlier in the Coastal and US ecotypes in response to Cd. Pairwise comparisons revealed only a few significant differences for the individual parameters (Table S2).

**Figure 2.**
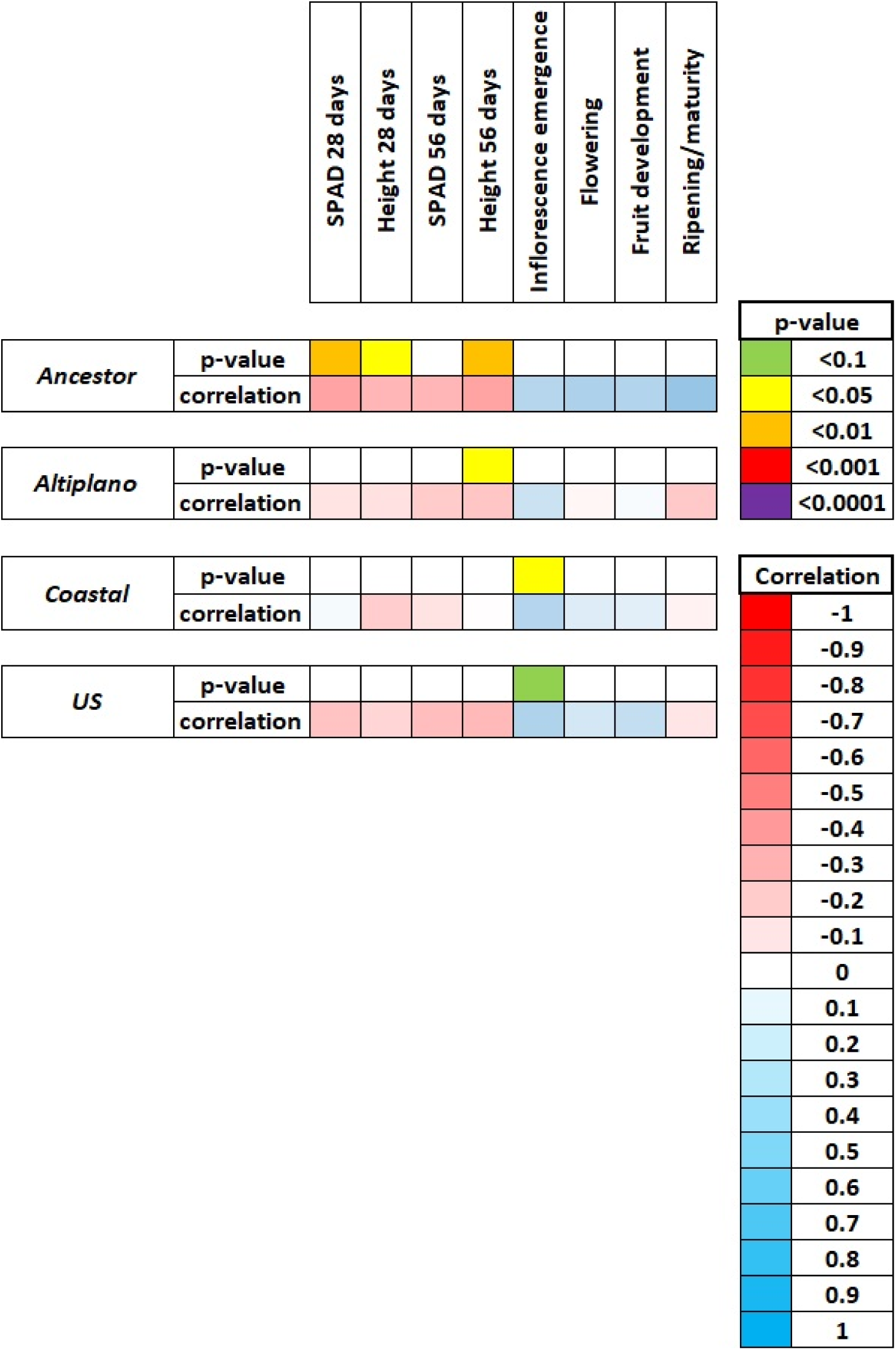
**Correlations between cadmium concentration in the growing media with early developmental parameters and phenological growth stages of four Chenopodium ecotypes**

### 3.2 Effect of Cd concentration and ecotype on root and shoot length and biomass at harvest

PERMANOVA results suggested that neither ecotype, Cd concentration, nor their interaction caused significant differences in plant length or biomass (all *p*>0.1). Cd concentration was also not significantly correlated with any of the plant growth measurements with the exception of the root:shoot length ratio, which was negatively correlated in the three *C. quinoa* ecotypes (Altiplano, Coastal and US), but not the Ancestor (Figure S1). With respect to pairwise differences, only the US ecotype was influenced by Cd concentration, with a lower root:shoot length ratio and higher root:shoot dry weight ratio with higher Cd concentration (Table S3). There were significant differences in seed yield among the ecotypes subjected to different levels of Cd (Table S3). Specifically, the Coastal and U.S. ecotypes produced significantly more seed than the Ancestor and Altiplano ecotypes regardless of Cd concentration, and at 5 mg/Kg Cd, the Ancestor outyielded the Altiplano. When comparing the impact of Cd on seed yield within each ecotype, the highest yield was obtained with plants grown at 5 mg/Kg Cd in the Ancestor, Coastal and U.S. ecotypes, and with 10 mg/Kg Cd in the Altiplano ecotype.

### 3.3 Concentrations of macronutrients, micronutrients and Cd in the harvested seeds

Results of the one-way ANOVA and subsequent Tukey post-hoc analysis of Cd concentrations in the grains conducted using ICP-MS analysis confirmed that there were significant differences in Cd accumulation among the individual genotypes and four ecotypes (Figure 3). In particular, the Coastal and US ecotypes had significantly greater concentrations of Cd than the Ancestor and Altiplano ecotypes. Results of the PERMANOVA analysis of Cd concentrations using ICP-OES further confirmed that ecotype (*F*= 41.903, *p*= 0.0001), potting media Cd concentration (*F*= 6.1953, *p*= 0.0037), and their interaction (*F*= 3.496, *p*= 0.0001) significantly affected Cd bioaccumulation in the grains. Cd accumulation in all four ecotypes was low when grown under 0 mg/Kg Cd, and differences in Cd accumulation among the four ecotypes followed a similar trend observed using ICP-MS (Table S4). Neither analysis indicated that potential differences in saponin concentrations in the grains among the genotypes were correlated with differences in Cd accumulation.

**Figure 3.**
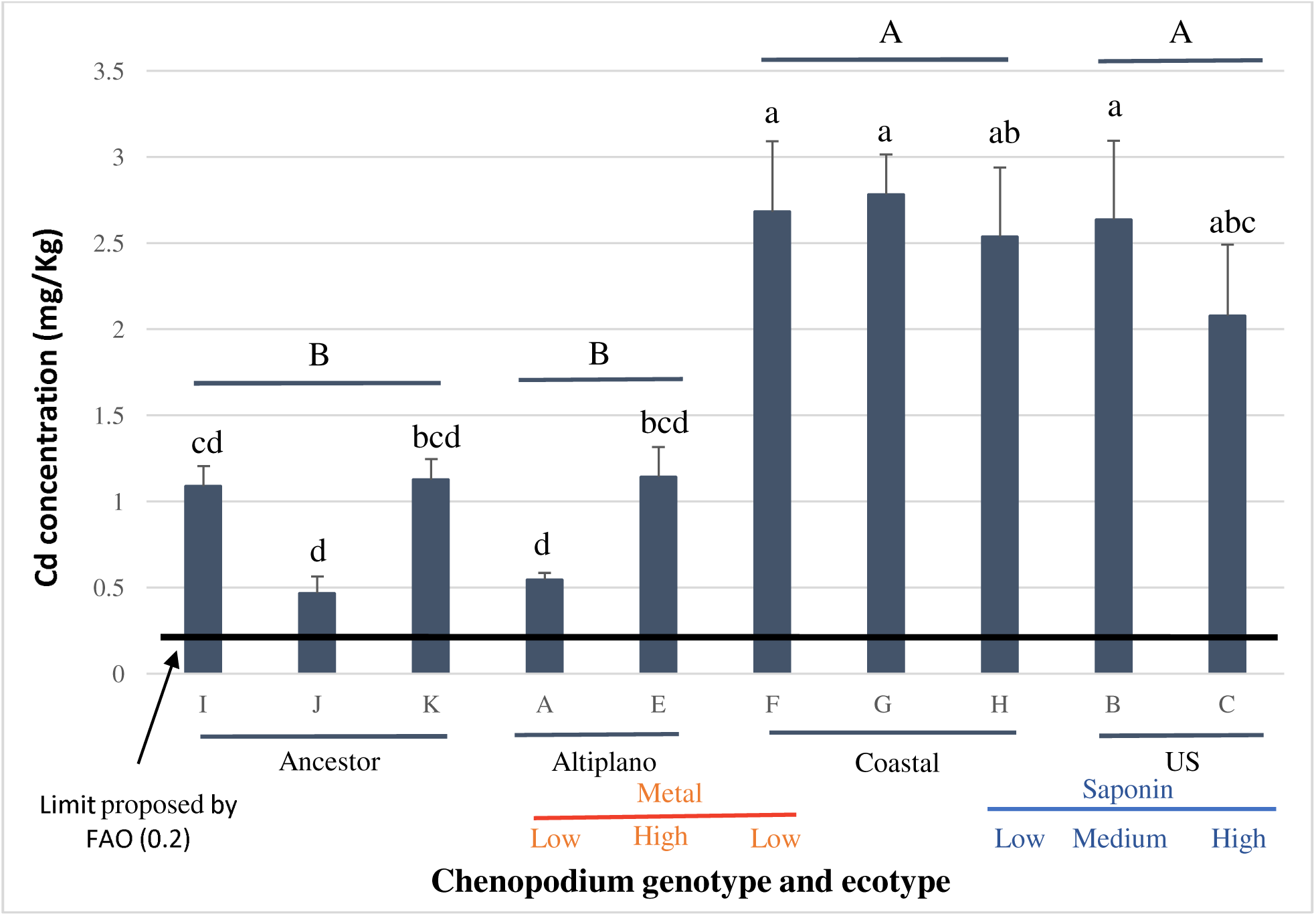
Cadmium (Cd) concentration in the grains of 10 Chenopodium genotypes (I, J, K, A, E, F, G, H, B, C) grouped into one of four ecotypes (ancestor, altiplano, coastal, US) grown in potting media containing 10 mg/Kg Cd. Genotypes that share the same lowercase letter and ecotypes that share the same capital letters are not significantly different (p<0.05). Genotypes with low or high metal denote previous findings by Bargava et al. (2008) for Cd uptake, and genotypes with low, medium and high saponin denote previous findings by Medina-Meza et al. (2008) for saponin concentrations in grains.

Correlations between the concentrations of macro and micronutrients in the grains with Cd concentrations in the potting media varied among the four Chenopodium ecotypes (Figure 4). The Ancestor ecotype had significantly lower concentrations of Ca, S, Cu, Ni and Si, and significantly higher concentrations of Mg. In contrast, the Altiplano ecotype had significantly higher concentrations of Ca, K and Mg, and all of the micronutrients; the Coastal ecotype had significantly higher concentrations of P, S, Fe and Zn; and the US ecotype had significantly higher concentrations of K, Mg, Fe, Na and Ni. Similar patterns were observed with respect to differences among the ecotypes and correlations between Cd and macro and micronutrients within the grains with a few exceptions (Figure 4). In the Ancestor ecotype, Mn was positively correlated with Cd, and in the US ecotype, Ca was positively correlated with Cd.

**Figure 4.**
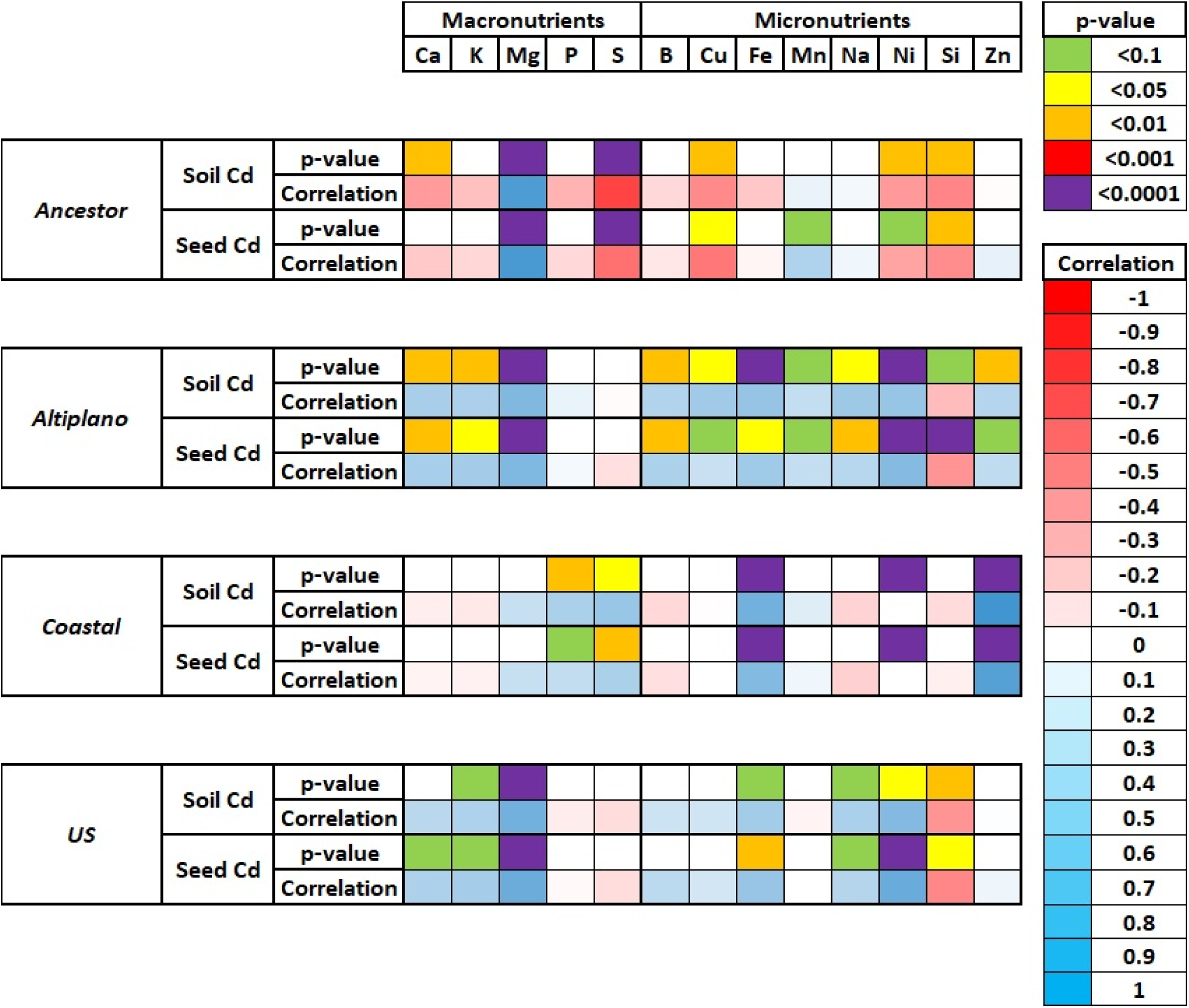
**Correlation between Cd in the growing media and grains of four Chenopodium ecotypes with the macro and micronutrients of the grains.**

**Figure 5.**
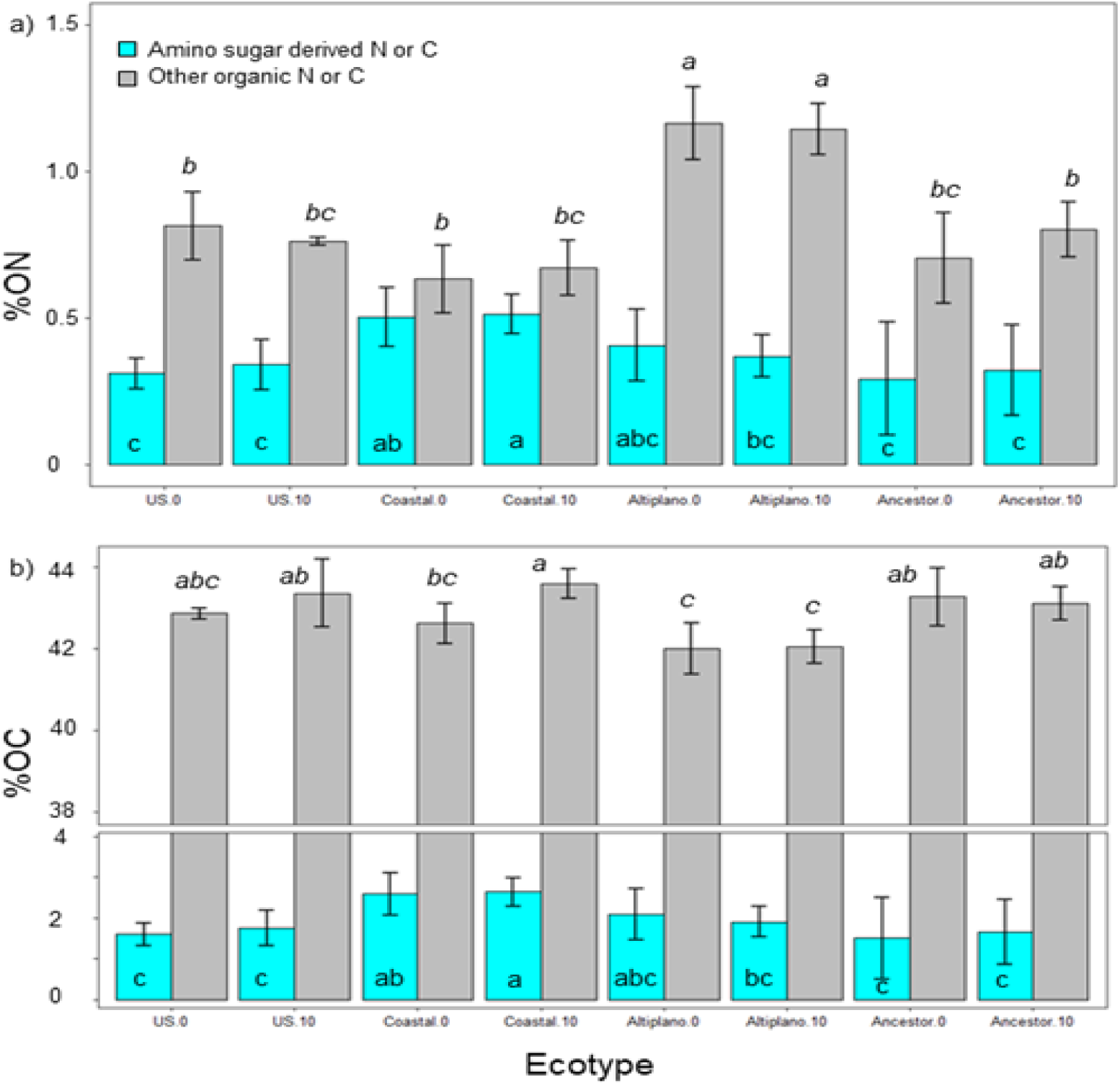
a) Percent contribution of amino sugar (AS) derived N (cyan bars) and other ON form (grey bars) to the ON content in the roots of four Chenopodium ecotypes. Bars sharing the same letter are not significantly different (for AS-N and other ON forms, P< 0.05; LSD). b) Percent contribution of AS derived C (cyan bars) and other OC form (grey bars) to the OC content in quinoa roots. Bars sharing the same letter are not significantly different (for AS-C, P< 0.05, LSD; for other OC forms, P< 0.01, LSD). The total ON and OC content is the sum of the AS- N and other ON forms, and the sum of the AS-C and other OC forms, respectively. In the name of the ecotypes 0= 0 ppm Cd, and 10= 10 ppm Cd

### 3.4 Carbon and nitrogen content in roots

Concentration of root nitrogen in all *Chenopodium* ecotypes ranged from 0.9% to 1.6% (Figure 6). Among these, the Altiplano ecotype exhibited the highest root nitrogen content at 1.6%, while the other ecotypes ranged from 0.9% to 1.1%. Conversely, root carbon content showed a narrower range, approximately 43% in the Ancestor and U.S. ecotypes and around 41% in the Altiplano and Coastal ecotypes. As a result, the mean C:N ratio differed among plants, following the order: Ancestor < U.S. ≈ Coastal < Altiplano (Table 2). In addition, isotope analysis of the samples showed that the δ^15^N values ranged from -0.9‰ to -0.4‰, while δ^13^C values ranged from -32.4‰ to -29.6‰ in roots from all ecotypes (Table 2). Across ecotypes, δ¹³C values were negatively correlated with the C:N ratio, following the order: Ancestor ≈ U.S. < Coastal < Altiplano (Figure 6). Notably the Ancestor ecotype was the most depleted in both ^13^C and ¹ N distinguishing it from the rest of the quinoa plants analyzed. Although ecotypic differences in C and N content as well as isotopic composition were observed, the Cd treatment had no significant effect on these traits, suggesting that the variations are primarily driven by ecotypic factors rather than Cd exposure.

**Figure 6.**
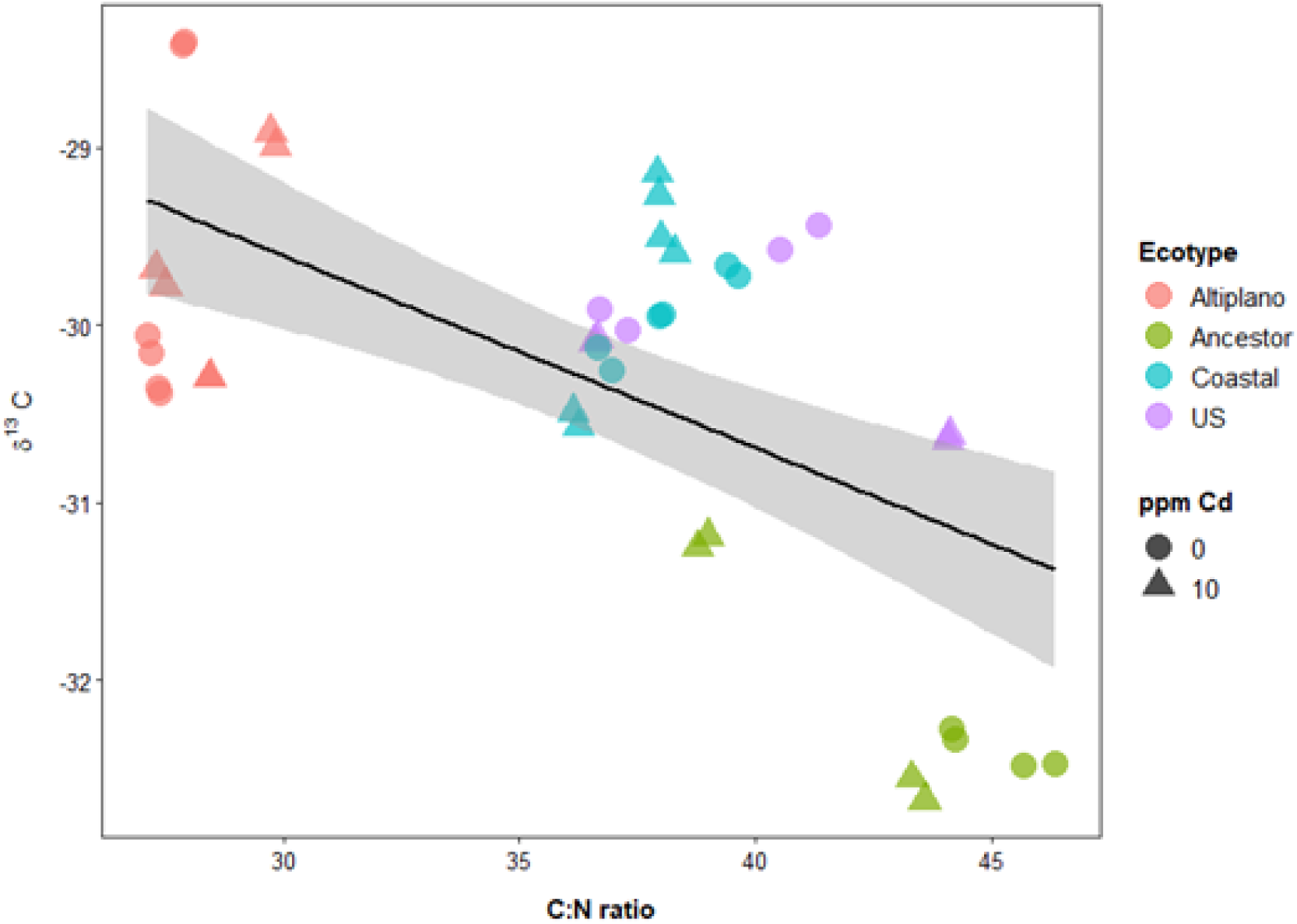
Correlation between the C:N ratio and δ^13^C (‰) values in the roots of four Chenopodium ecotypes

**Table 2.**
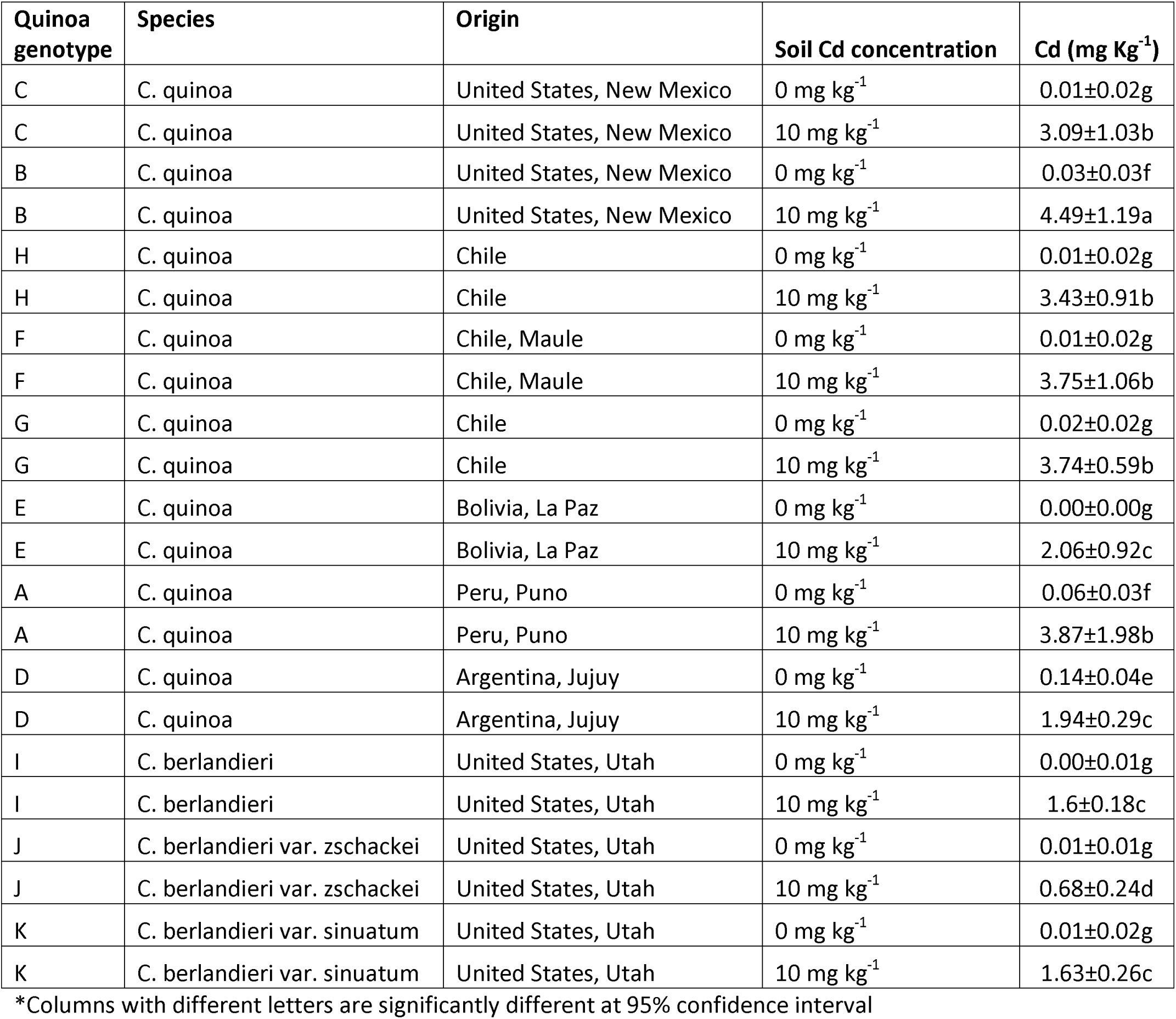
Cd concentration in seeds of the different quinoa genotypes

Table – Cd concentration in seeds of the different quinoa genotypes
| Quinoa genotype | Species | Origin | Soil Cd concentration | Cd (mg Kg<sup>-1</sup>) |
| --- | --- | --- | --- | --- |
| C | C. quinoa | United States, New Mexico | 0 mg kg<sup>-1</sup> | 0.01±0.02g |
| C | C. quinoa | United States, New Mexico | 10 mg kg<sup>-1</sup> | 3.09±1.03b |
| B | C. quinoa | United States, New Mexico | 0 mg kg<sup>-1</sup> | 0.03±0.03f |
| B | C. quinoa | United States, New Mexico | 10 mg kg<sup>-1</sup> | 4.49±1.19a |
| H | C. quinoa | Chile | 0 mg kg<sup>-1</sup> | 0.01±0.02g |
| H | C. quinoa | Chile | 10 mg kg<sup>-1</sup> | 3.43±0.91b |
| F | C. quinoa | Chile, Maule | 0 mg kg<sup>-1</sup> | 0.01±0.02g |
| F | C. quinoa | Chile, Maule | 10 mg kg<sup>-1</sup> | 3.75±1.06b |
| G | C. quinoa | Chile | 0 mg kg<sup>-1</sup> | 0.02±0.02g |
| G | C. quinoa | Chile | 10 mg kg<sup>-1</sup> | 3.74±0.59b |
| E | C. quinoa | Bolivia, La Paz | 0 mg kg<sup>-1</sup> | 0.00±0.00g |
| E | C. quinoa | Bolivia, La Paz | 10 mg kg<sup>-1</sup> | 2.06±0.92c |
| A | C. quinoa | Peru, Puno | 0 mg kg<sup>-1</sup> | 0.06±0.03f |
| A | C. quinoa | Peru, Puno | 10 mg kg<sup>-1</sup> | 3.87±1.98b |
| D | C. quinoa | Argentina, Jujuy | 0 mg kg<sup>-1</sup> | 0.14±0.04e |
| D | C. quinoa | Argentina, Jujuy | 10 mg kg<sup>-1</sup> | 1.94±0.29c |
| I | C. berlandieri | United States, Utah | 0 mg kg<sup>-1</sup> | 0.00±0.01g |
| I | C. berlandieri | United States, Utah | 10 mg kg<sup>-1</sup> | 1.6±0.18c |
| J | C. berlandieri var. zschackei | United States, Utah | 0 mg kg<sup>-1</sup> | 0.01±0.01g |
| J | C. berlandieri var. zschackei | United States, Utah | 10 mg kg<sup>-1</sup> | 0.68±0.24d |
| K | C. berlandieri var. sinuatum | United States, Utah | 0 mg kg<sup>-1</sup> | 0.01±0.02g |
| K | C. berlandieri var. sinuatum | United States, Utah | 10 mg kg<sup>-1</sup> | 1.63±0.26c |

\*Columns with different letters are significantly different at 95% confidence interval

Fungal-derived amino sugars (GlcN, GalN) were predominant in the root tissues, while ManN and MurN were not detected (Table 3). The Coastal ecotype exhibited the highest contribution of amino sugar derived nitrogen, accounting for 44% of total root nitrogen, whereas this contribution was lower in other ecotypes (19% in Ancestors, 25% in Altiplano, and 29% in US). Similarly, the amino sugar derived carbon was more abundant in the Coastal (5.9%) and Altiplano (4.6%) ecotypes compared to the US (3.8%) and the Ancestor (2.3%) (Figure 6).

The correlation between microbial amino sugars and Cd in the potting media, along with plant phenological traits and elemental concentrations in quinoa seeds, highlights distinct patterns across all the ecotypes (Figure S2). Specifically, Cd in the potting media was negatively correlated with the Coastal ecotype and positively correlated with the U.S. ecotype. In the Altiplano ecotype, Cd showed positive correlations with inflorescence emergence and seed yield, while negative correlations were observed with flowering, fruit development, and ripening.

There were also distinct differences in how Cd in the potting media influenced elemental concentrations in the seeds. In the Ancestor ecotype, Cd concentrations were positively correlated with glucosamine-derived nitrogen (GlcN-N) and glucosamine-derived carbon (GlcN- C). Similarly, all microbial amino sugars were positively correlated with Cd in the Altiplano and U.S. ecotypes, whereas in the Coastal ecotype, they showed a negative correlation. Similarly, there were distinct differences in the concentrations of other elements in the seeds that are often influenced by the presence of Cd due to changes in the activity of root transporters. For example, Fe concentration was positively correlated with Cd concentrations in the seeds of the Altiplano ecotype, negatively correlated with the Coastal ecotype, and there was no difference in the Ancestor or U.S. ecotypes. For Zn concentration, both the Ancestors and Altiplano ecotypes were positively correlated with Cd, the coastal ecotype had a negative correlation, and there was no difference in the U.S. ecotype. Finally, Na concentration was negatively correlated in the Ancestor ecotype, and positively correlated with the U.S. ecotype.

## 4 Discussion

### 4.1 Cd bioaccumulation varies based on Chenopodium genotype and ecotype

Results of this study confirm previous reports (Bhargava et al., 2008) showing that quinoa genotypes differ in their capacity to uptake and accumulate Cd in their grains or seeds at harvest (Figure 3). Moreover, our study indicates that domestication and the region where the genotypes were selected is important in differentiating uptake among quinoa genotypes, with genotypes selected near coastal regions of South America and recent selections in the U.S. having greater uptake than the Altiplano or wild Ancestor ecotypes (Figure 1). To our knowledge, this is the first time this has been reported.

Since the germplasm included in this study was narrow, differences in Cd accumulation may be due to the parent materials used during selection within these regions. For example, similar findings have been reported in maize (*Zea mays*), where differences in Cd uptake were observed among genotypes within the same domestication lineage (Vielle-Calzada et al., 2009). In that study, chromosomal regions of low nucleotide diversity containing genes involved in heavy-metal detoxification were observed, indicating that environmental changes may have been important in selecting for higher uptake. Similar results were observed in durum wheat (*Triticum durum*), where domesticated lines exhibited higher Cd uptake and the presence of specific root transporters that facilitated Cd absorption compared to their undomesticated relatives (Macaferri et al., 2019). Thus, domestication and selection of quinoa and other crops in environments with higher concentrations of toxic heavy metals like Cd in soil may induce adaptions within plants to help them tolerate, and possibly even benefit from, higher concentrations of these metals in their tissues. For example, Poschenrider et al., 2006 theorized and found evidence to suggest that some plants have evolved the capacity to withstand heavy metal stress and tolerate higher levels of metals in their vegetative tissues as a defense mechanism against fungal pathogens and herbivore pests. However, this accumulation can pose risks to human health when these plant tissues are consumed, highlighting the importance of identifying genotypes that limit the uptake of toxic heavy metals like Cd to ensure food safety. In this study, both the wild crop ancestors and quinoa genotypes selected in the Altiplano region had lower Cd uptake relative to the other genotypes, indicating that these lines could serve as germplasm for the development of improved quinoa cultivars with low uptake. Understanding the mechanisms mediating these differences will provide important clues to help select for this important trait.

While our results were consistent with a previous study by Bargava et al. (2008) who compared Cd uptake differences among some of the same genotypes, we also observed distinct differences from their findings. For example, genotype F (PI 614886) had greater Cd uptake in our study relative to the other genotypes, while it had lower uptake relative to the others in the study by Bargava et al. (2008). This could be due to differences in the growth media used in the two experiments. A wide range of soil properties including pH, the types of clays and organic materials present, concentrations of other nutrients such as phosphorous and microbiome composition can all alter the bioavailability of heavy metals like Cd in soil (Shahid et al., 2017; Zea et al., 2022; Souza et al., 2022). Plant roots release organic substances (exudates) that can further alter the bioavailability of heavy metals in soil directly (Mench et al., 1991), or indirectly via modification of soil and root microbiomes (Caracciolo and Terenzi, 2021). The composition of root exudates has been shown to vary among plant genotypes (Micallef et al., 2009), which could be an important factor regulating differences in Cd uptake among quinoa genotypes.

However, since soil is generally a greater factor in shaping the composition and activity of root microbiomes (Abdelrazek et al., 2020a; 2020b), plant genotypes could still respond differently when grown in different types of soils and other substrates. Consequently, future studies investigating how different quinoa genotypes interact with different soil types and their microbiomes are needed to select for lower uptake in breeding programs and ensure that results will be heritable under field conditions.

Like many plants, quinoa tissues are vulnerable to attacks by a diverse assortment of pathogens and insect pests. One defense strategy used by some plants against these pests is the production of saponins, which are biologically active toxic compounds (Zaynab et al., 2021). Quinoa plants produce significant amounts of these compounds in their pericarps and seed coats. Since saponins also impart a bitter flavor during cooking, the identification of quinoa germplasm and development of new cultivars with lower saponin content has been a major goal in breeding programs (Zurita-Silva et al., 2014). Saponins can also bind heavy metals like Cd (Nobahar et al., 2021), so we theorized that genotypes with different levels of saponins in their seeds might also influence Cd accumulation. To test this hypothesis, we compared Cd uptake in three quinoa genotypes previously shown to differ in the concentrations of saponins in their seeds (Table 1; Medina-Meza et al., 2016). We did not observe significant differences in Cd accumulation among these three genotypes (Fig. 3), which indicates that saponins do not appear to play a role in Cd accumulation. However, we did not explicitly measure saponin concentrations, so it is possible that this trait may not have been expressed under our growing conditions. Other recent studies have also demonstrated that while light abrasive decortication to remove seed coats does help in reducing lead and arsenic contamination in quinoa grains, it does not help in removing Cd (Roman-Ochoa et al., 2023), providing further evidence that seed coat traits may not help in reducing this food safety risk.

### 4.2 Quinoa ecotypes differ in phenological development in response to Cd

Quinoa genotypes from the Ancestor and Altiplano ecotypes appeared to be more adversely affected by the presence of Cd in this study (Figure 2), which may have contributed to their lower Cd uptake (Figure 3). Cd is generally detrimental for plant growth, causing impairments in several plant metabolic pathways leading to inhibition of photosynthesis and/or the fixation and assimilation of important nutrients like nitrogen (Sebastian & Prasad, 2015a; 2015b). Cd can also negatively affect stomatal opening and transpiration processes (Chandra & Kang, 2016), leading to lower plant biomass. In response, some plants employ strategies to restrict the uptake of toxic heavy metals like Cd from soils or potting media and reduce subsequent translocation to aboveground tissues where they can negatively affect critical plant processes (Khanna et al., 2022). For example, the production molecules such as phytochelatins and glutathione have been shown to restrict translocation of Cd to carrot leaves plants by promoting vacuole sequestration in roots (Gao et al., 2022). Plants can also host specific microbes within their root systems such as dark septate endophytes that reduce Cd uptake by altering root morphology and root cellular structures, binding Cd in roots and fungal hyphae and thereby restricting translocation to above ground tissues (Shen et al., 2020; Chen et al., 2023). Like other plant traits, differences in the capacity to host and benefit from root-associated microbes can vary among plant genotypes (Jaiswal et al., 2020; Trivino et al., 2023), and is another factor that should be explored in future studies to determine why these quinoa genotypes vary in Cd uptake.

Although the coastal and US ecotypes did not exhibit reduced growth in response to Cd stress (Figure 2), inflorescence emergence occurred earlier, indicating stress, though less pronounced than in the Ancestor and Altiplano genotypes. Similar responses have been observed in *Arabidopsis thaliana*, where plants accelerated inflorescence emergence in response to Cd stress (Keunen et al., 2011). All genotypes with the exception of the Ancestors exhibited a negative correlation (p<0.1) between their root:shoot length ratio with Cd concentration in the potting media. Specifically, the plants had shorter roots and longer shoots in response to the Cd. In contrast, many other plant species (Rolón-Cárdenas et al., 2022), including *Triticum aestivum* and *Arabidopsis thaliana* (Zhang et al., 2022; Sofo et al., 2022), often have both smaller roots and shoots in response to Cd. This may be one of the ways quinoa plants use to tolerate Cd stress and promote higher uptake of this compound. For example, some plants are able to mitigate heavy metal stress by upregulating the expression of phytohormones like salicylic acid and gibberellin, which can in turn alter the expression of various transporters as well as other biochemical reactions like ROS scavenging that can help protect plants from heavy metal stress (Emamverdian et al., 2020). Upregulation of these phytohormones can also increase plant growth and may contribute to the hormetic response often observed when some plants are exposed to low levels of Cd stress (Zea et al., 2022). This also may have contributed to the plant growth responses we observed in quinoa, such as greater seed yield in the Ancestor, Coastal and U.S. genotypes when subject to 5 mg/Kg Cd, and in the Altiplano at 10 mg/Kg (Figure S2).

Interestingly, differences in uptake among some plant genotypes can be due to a dilution factor, with greater yield being correlated with lower Cd uptake, but that was not the case in this study. Instead, we observed both greater seed yield (Figure S2) and Cd accumulation in the Coastal and U.S. ecotypes (Figure 3). Again, this indicates that these ecotypes have developed adaptations that allow them to thrive in areas with high concentrations of soil Cd.

### 4.3 Differences in elemental profiles among the seed of different ecotypes in response to Cd

The presence of toxic heavy metals like Cd in soils and other growing media are well known for their potential to interfere with elemental homeostasis in plants. For example, since some root transporters cannot differentiate between Fe and Cd (Kar et al., 2021; Liu et al., 2022), some plants will shut down these transporters to reduce Cd uptake, in some cases resulting in Fe-induced chlorosis (Leskova et al., 2017). Similarly, root transporters within the ZIP (Zrt/Irt-like protein) family have been shown to control both Cd and Zn uptake, which can also have important implications for Zn uptake (Grotz et al., 2006). The presence of greater concentrations of plant available Fe and Zn in the soil can also reduce Cd stress and uptake by competing for the same transport pathways (Rizwan et al., 2019). In this study, all of the quinoa ecotypes other than the Ancestor had greater concentrations of Fe in response to Cd, and the Altiplano and Coastal genotypes had greater concentrations of Zn (Figure 4). Consequently, quinoa plants may have evolved mechanisms over the course of domestication to prevent downregulation of these transporters as part of the plant’s capacity to accommodate Cd uptake. Root interaction with microbes that can regulate Fe and Zn availability in soil could also have influenced these dynamics Na is another element often shown to influence Cd uptake in crops, but whether this increases or decreases Cd uptake depends on the crop and even cultivar (Cheng et al., 2018). The mechanisms regulating how plants control Cd uptake in response to Na include changes in the production of antioxidants and other metabolites that can act as ligands, facilitating translocation of Cd within plants, reduce symptoms of stress, and/or influence root transporters (Mei et al., 2014; Cheng et al., 2018). Interestingly in a study with amaranth (*Amaranthus mangostanus* L.), another member of the *Chenopodium* family, Na reduced Cd uptake in a salt-sensitive cultivar, and increased uptake in a salt-tolerant cultivar (Mei et al., 2014), whereas the opposite trend has been observed in wheat and maize (Muhling and Lauchli, 2003; Sepehr and Ghorbanli, 2006). In one quinoa study using the cultivar ‘Puno’, the application of Na reduced Cd uptake (Abdal et al., 2021), indicating a positive impact in quinoa. In our experiment, we observed greater grain concentrations of Na in response to Cd in both the Altiplano and US ecotypes (Figure 4), indicating that Na may be an important moderator of Cd uptake in these ecotypes. However, since these two ecotypes responded differently with respect to the amount of Cd they took up, the ecotypes may be responding with contrasting mechanisms.

Two additional elements that are known to influence Cd-induced plant stress and uptake are Mg and Ca. Similar to Na, both elements can indirectly influence Cd uptake by modifying plant stress responses, and/or directly affect uptake via interference with root transporters, thereby increasing or decreasing Cd uptake (Kikuchi et al., 2008; Mleczeck et al., 2011; Zhang et al., 2020). In our study, all ecotypes, except for the Coastal, had greater concentrations of Mg in their grains in response to Cd, whereas the Altiplano and US ecotypes had greater concentrations of Ca in response to Cd and the opposite was observed for the Ancestor (Figure 4). Given the differing responses of these ecotypes in terms of Cd uptake, it is challenging to hypothesize how these elements might be influencing physiological functions across this diverse group of plants to regulate Cd uptake. It is interesting to note however, that the Coastal ecotype, which theoretically descends from the Ancestor and then Altiplano ecotype h, exhibited high uptake of Cd, but did not exhibit any Ca or Mg-induced changes in response to Cd. Thus, further insights comparing how these elements influence Cd uptake in genotypes representing the Altiplano and Coastal would be interesting to explore.

### 4.4 C and N content in the roots of the Chenopodium ecotypes

Quinoa is a C3 plant, resistant to several adverse abiotic stress factors including drought, frost, and salinity (Jacobsen et al., 2003). N is a critical determinant in all plant developmental stages, from seed germination to senescence, and this element is generally a key factor limiting crop yield and quality (Erley Schulte auf’m et al., 2005). In this study, all pots received the same amount of N fertilizer, thus the difference in N concentration and isotope values observed could be attributed to the Cd content in the growing media, and/or the quinoa ecotype/genotype. N content in roots (1.2%, on average) was within the range of previous studies in growing mediums with no additional N input (Kakabouki et al., 2013), and was comparable to the roots of legumes and grasses (Dijkstra et al., 2003; Robinson et al., 2000). Root C content (43%, on average) was also within the expected value for herbs, but close to woody root values (Ma et al., 2018).

The C:N ratio is a commonly used indicator of plant growth and quality. Paired *t*-tests were conducted to evaluate the effect of Cd on this ratio by comparing plants grown in Cd-amended versus Cd-free media. Our results showed that Ancestor plants grown in Cd-amended media had a significantly lower C:N ratio compared to those grown in Cd-free conditions. In contrast, other ecotypes did not exhibit significant differences between Cd treatments. Among ecotypes, the Altiplano roots were distinct, displaying the lowest C:N ratio, suggesting that the Altiplano ecotype may share characteristics more similar to the Ancestor.

### 4.5 Fungal sources are the primary contributors of amino sugar derived C and N in Chenopodium roots

Quinoa hosts a diverse community of root-associated fungi, acquired either through vertical transmission via seed or horizontal recruitment from the soil in response to root exudates. These fungal associations play a critical role in nutrient acquisition and enhancing plant resilience to biotic and abiotic stresses (Reeve et al., 2016; Triviño et al., 2023). For example, most plants associate with arbuscular mycorrhizal fungi (AMF), which are well known for their capacity to promote plant growth by providing multiple benefits including increasing P, Zn, and Fe uptake. While quinoa is generally considered non-mycorrhizal due to its classification within the *Amaranthaceae* family (Urcelay et al., 2011), phospholipid fatty acid analyses suggest that it can exhibit mycotrophic traits and support moderate levels of AMF (Vestberg et al., 2012). Beyond AMF, quinoa roots also host many other genera including *Penicillium*, *Phoma*, and *Fusarium* (Hinojosa et al., 2018). Previous studies have shown that *C. quinoa* establishes symbiotic associations with root endophytic fungi, enhancing its morphological and physiological responses to abiotic stresses such as drought and salinity. These symbioses promote the secretion of antioxidant compounds under stress conditions, helping to counteract salinity-induced oxidative stress (González-Teuber et al., 2022). Consistently, our study found that the Coastal ecotype exhibited a substantially higher contribution of amino sugar-derived N and C in its roots, primarily GlcN and GalN (Table 3). This may be linked to the potential role of fungi in enhancing the salinity tolerance of the plants, as coastal soils are typically more saline. The Coastal ecotype did not exhibit greater Na uptake in response to Cd (Figure 4), providing further evidence that these ecotypes are more salt-tolerant.

Interestingly, amino sugars in the quinoa roots were negatively correlated to Cd, Fe and Zn concentrations in the seed from the Coastal genotypes (Figure S2), which had the highest Cd uptake (Figure 3). In contrast, these elements were positively correlated in the Altiplano and Ancestor ecotypes, which had much lower Cd uptake. This indicates that the capacity of quinoa genotypes to recruit and host fungi that can regulate the bioavailability and uptake of these critical elements could vary. For example, several studies have demonstrated that fungal biomass from several species is capable of immobilizing Cd in wastewater (Tsekova et al., 2010; Cai et al., 2016; Ali et al., 2021). Future studies aimed at identifying the fungal species associated with these differences would be highly beneficial to understanding how they can influence Cd dynamics in quinoa roots.

### 4.6 Cadmium does not influence the C and N content in quinoa roots

Cadmium concentrations in the soil did not induce significant alterations in root N or root C content, whether expressed as bulk percentages or as amino sugar-derived elements, across most ecotypes. However, the Coastal ecotype showed a significant increase in root carbon C when cultivated in Cd-amended soil, suggesting a distinct physiological response to Cd exposure.

Previous studies (Robinson et al., 2000; Yousfi et al., 2012) have used δ^13^C and δ^15^N values as indicators of plant responses to environmental stresses such as salinity, drought, and N limitation. Our results indicate that the Coastal ecotype displayed higher δ^13^C and δ^15^N values specially compared to those of the Ancestor ecotype, suggesting greater tolerance to environmental stresses. Given the variation in Cd uptake across quinoa genotypes, the effect of Cd and other toxic heavy metals on the isotopic values of quinoa roots require further analysis (Fig. S3).

## 5. Conclusions and future research

Once toxic heavy metals like Cd become enriched in farm soils, remediation strategies are limited, so developing new improved quinoa cultivars that can restrict uptake while maintaining other important agronomic and end-use quality traits is critical for addressing food safety concerns. Our study provides further evidence to support genetic differences in Cd uptake among quinoa germplasm indicating that it will be possible to select for this important trait in breeding programs. Like other plants, the potential to tolerate and accumulate greater concentrations of Cd appears to have increased along domestication, and this may provide some benefits with respect to tolerating various biotic and abiotic stress factors. Understanding these trade-offs to selecting for low Cd uptake will be critical for developing improved varieties that retain other valuable agronomic traits. Since using wild crop ancestors in breeding programs is challenging due to linkage drag associated with non-desirable traits, using landraces and other modern germplasm may be a better approach. In our study, there are clear differences between landrace and modern genotypes representing the Altiplano and Coastal ecotypes with respect to Cd uptake and resilience against other potential stress responses. Consequently, future studies that include more genotypes from these two ecotypes could provide valuable insights into the mechanisms regulating how these ecotypes respond to environmental stress and Cd.

