## Supplementary materials for "Variability in cadmium accumulation in quinoa grain: potential role of crop domestication and other factors mediating differences in resilience and uptake"

**Table S1. The regression (R) value, limit of detection (LOD) and limit of quantification (LOQ) used to quantify the concentrations of several elements in quinoa seed.**

| **Element** | **R** | **LOD (**µg per kg ^-1^**)** | **LOQ (**µg per kg ^-1^**)** |
| --- | --- | --- | --- |
| Boron | 0.9702 | 21.2198 | 70.7327 |
| Calcium | 0.9402 | 0.9585 | 3.1950 |
| Cadmium | 0.9998 | 0.0686 | 0.2289 |
| Cobalt | 0.9998 | 0.1950 | 0.6499 |
| Copper | 0.9934 | 0.5509 | 1.8362 |
| Iron | 0.9977 | 0.6876 | 2.2920 |
| Potassium | 0.9626 | 19.8812 | 66.2708 |
| Magnesium | 0.9992 | 0.2017 | 0.6722 |
| Manganese | 0.9995 | 0.0216 | 0.0701 |
| Sodium | 0.9920 | 2.4014 | 8.0045 |
| Nickel | 0.9997 | 0.3189 | 1.0631 |
| Phosphorous | 0.9987 | 3.9244 | 13.0812 |
| Sulfur | 0.9922 | 5.9408 | 19.8027 |
| Silicon | 0.9976 | 0.5114 | 1.7047 |
| Zinc | 1.0000 | 0.0197 | 0.0657 |

**Table S2. Differences in phenological development parameters of Chenopodium ecotypes grown in soil containing 0 and 10 mg/Kg cadmium (Cd)**

|  |  | **Phenological development** | | | | **Plant height (cm)** | | **SPAD meter readings** | | |
| --- | --- | --- | --- | --- | --- | --- | --- | --- | --- | --- |
| **Quinoa ecotype** | **Soil Cd concentration** | **Inflorescence emergence** | **Flowering** | **Fruit development** | **Ripening** | **28 days after planting** | **56 days after planting** | **28 days after planting** | **56 days after planting** | **Harvest** |
| Ancestor | 0 mg/Kg | 34±90.78 a | 47.45±11.98 a | 58.55±11.39 a | 75.36±13.43 b | 18.13±10.38 a | 105.83±19.14 a | 62.64±11.61 b | 45.06±6.31 a | 34.00±9.78 a |
| Ancestor | 5 mg/Kg | 37.17±70 a | 51.25±9.75 a | 62.75±12.29 a | 83.08±11.73 a | 11.04±4.75 a | 105.67±7.00 a | 72.38±5.44 a | 36.10±6.60 a | 37.17±7.00 a |
| Ancestor | 10 mg/Kg | 36.82±9.50 a | 53.55±11.36 a | 65.09±13.42 a | 84.73±12.91 a | 10.67±9.30 a | 82.45±12.73 b | 63.93±10.03 b | 39.76±7.30 a | 36.82±9.50 a |
| Altiplano | 0 mg/Kg | 33.94±5.61 a | 62.57±13.64 a | 74.92±8.77 a | 125.89±11.12 a | 17.39±7.20 a | 104.56±23.94 a | 66.84±6.57 a | 34.22±4.84 a | 33.94±5.61 a |
| Altiplano | 5 mg/Kg | 34.59±6.32 a | 59.36±10.77 a | 68.27±8.76 a | 125.40±9.43 a | 17.89±6.39 a | 98.61±11.87 ab | 63.17±9.27 a | 36.91±7.50 a | 34.59±6.32 a |
| Altiplano | 10 mg/Kg | 35.94±5.99 a | 62.92±19.33 a | 72.73±10.03 a | 126.10±17.16 a | 15.11±6.86 a | 88.17±18.10 b | 64.68±6.15 a | 32.89±5.16 a | 35.94±5.99 a |
| Coastal | 0 mg/Kg | 29.56±2.06 ab | 41.33±2.43 a | 53.50±1.82 a | 83.00±4.12 a | 28.28±4.78 a | 109.00±13.79 a | 67.21±6.16 a | 44.16±5.17 a | 29.56±2.06 ab |
| Coastal | 5 mg/Kg | 29.78±1.26 ab | 41.22±2.46 a | 53.06±2.26 a | 81.72±1.90 a | 27.56±4.89 a | 91.11±11.11 b | 63.65±6.37 a | 45.12±7.56 a | 29.78±1.26 ab |
| Coastal | 10 mg/Kg | 30.88±2.06 a | 42.35±2.64 a | 54.29±2.39 a | 82.65±3.76 a | 25.97±4.51 a | 79.78±28.00 b | 59.23±6.83 a | 44.40±4.20 a | 30.88±2.06 a |
| US | 0 mg/Kg | 33.64±2.91 a | 49.09±4.21 a | 59.27±4.47 b | 103.82±17.4 a | 23.17±8.61 a | 74.50±31.75 b | 53.43±10.85 b | 43.98±4.65 a | 33.64±2.91 a |
| US | 5 mg/Kg | 34.55±3.36 a | 51.92±4.06 a | 65.83±8.41 a | 103.33±17.83 a | 22.50±5.39 a | 105.92±9.51 a | 64.67±5.32 a | 38.28±3.27 a | 34.55±3.36 a |
| US | 10 mg/Kg | 35.83±3.19 a | 50.58±3.85 a | 62.75±6.96 a | 103.82±19.59 a | 22.67±3.73 a | 75.00±16.26 b | 59.62±7.45 ab | 39.31±6.85 a | 35.83±3.19 a |

*Values within each column and genotype with the same lowercase letters are not significantly different at 95% confidence interval

**Table S3. Differences in root and shoot length, root and shoot wet and dry biomass, and their ratios of Chenopodium ecotypes grown in soil containing 0 and 10 mg/Kg cadmium (Cd)**

|  |  |  | **Biomass length (cm)** | | | **Wet biomass weight (g)** | | | **Dry biomass weight (g)** | | |
| --- | --- | --- | --- | --- | --- | --- | --- | --- | --- | --- | --- |
| **Quinoa ecotype** | **Soil Cd concentration** | **Seed yield (g)** | **Root** | **Shoot** | **Root:shoot ratio** | **Root** | **Shoot** | **Root:shoot ratio** | **Root** | **Shoot** | **Root:shoot ratio** |
| Ancestor | 0 mg/Kg | 6.96±5.32 e | 101.45±30.29 a | 38.00±18.43 a | 0.42±0.25 a | 109.35±59.97 a | 37.83±12.76 a | 0.44±0.23 a | 27.82±16.42 a | 8.91±5.32 a | 0.35±0.11 a |
| Ancestor | 5 mg/Kg | 24.96±9.17 c | 101.08±24.65 a | 31.50±6.9 a | 0.32±0.09 a | 128.53±59.43 a | 54.84±23.85 a | 0.62±0.55 a | 34.00±16.98 a | 10.92±6.4 a | 0.36±0.18 a |
| Ancestor | 10 mg/Kg | 6.68±5.22 e | 97.83±37.40 a | 33.33±13.49 a | 0.38±0.18 a | 121.21±68.77 a | 53.96±34.05 a | 0.50±0.23 a | 31.25±21.12 a | 13.09±13.22 a | 0.43±0.14 a |
| Altiplano | 0 mg/Kg | 4.59±7.79 e | 166.06±40.91 a | 38.15±10.39a | 0.25±0.08 ab | 231.42±156.73 a | 23.30±10.28 a | 0.18±0.14 a | 62.25±35.43 a | 7.79±5.47 a | 0.20±0.13 a |
| Altiplano | 5 mg/Kg | 12.85±15.26 de | 158.28±58.56 a | 41.86±17.01 a | 0.29±0.09 a | 196.13±139.86 a | 31.26±25.21 a | 0.31±0.31 a | 57.31±31.24 a | 6.86±4.45 a | 0.20±0.17 a |
| Altiplano | 10 mg/Kg | 3.12±4.79 e | 165.11±38.15 a | 32.5±7.92 a | 0.22±0.07 b | 247.56±153.45 a | 35.38±28.80 a | 0.21±0.13 a | 59.56±29.58 a | 11.14±7.01 a | 0.24±0.13 a |
| Coastal | 0 mg/Kg | 18.52±4.96 cd | 95.19±13.12 a | 35.06±7.28 a | 0.37±0.07 a | 62.86±26.01 a | 36.75±9.13 a | 0.64±0.18 b | 22.61±5.91 a | 7.33±2.00 a | 0.35±0.15 a |
| Coastal | 5 mg/Kg | 32.15±7.35 b | 93.25±11.35 a | 30.28±10.33 a | 0.31±0.10 b | 63.90±58.14 a | 46.53±25.49 a | 0.87±0.39 a | 23.44±12.04 a | 8.83±4.77 a | 0.39±0.16 a |
| Coastal | 10 mg/Kg | 15.13±6.51 d | 99.17±14.63 a | 32.50±11.7 a | 0.34±0.17 ab | 62.16±57.41 a | 32.63±15.79 a | 0.75±0.52 ab | 21.72±12.51 a | 6.39±2.93 a | 0.33±0.17 a |
| US | 0 mg/Kg | 24.22±16.87 c | 91.42±57.92 a | 31.82±9.95 a | 0.49±0.20 a | 94.22±79.01 a | 30.31±18.18 a | 0.41±0.26 a | 43.67±37.66 a | 9.09±6.16 a | 0.3±0.14 b |
| US | 5 mg/Kg | 49.33±11.01 a | 132.28±28.00 a | 38.04±9.92 a | 0.27±0.09 b | 80.67±44.25 a | 47.25±21.05 a | 0.69±0.34 a | 35.58±18.78 a | 16.83±13.04 a | 0.48±0.23 a |
| US | 10 mg/Kg | 7.8±12.41 d | 134.58±31.88 a | 33.42±8.93 a | 0.26±0.07 b | 77.27±70.37 a | 31.26±24.75 a | 0.58±0.38 a | 26.50±13.92 a | 9.25±5.24 a | 0.39±0.24 ab |

*Values within each column and ecotype with different letters are significantly different at 95% confidence interval

**Table S4. Elemental composition of cadmium and other elements in Chenopodium seeds grown in soil containing 0 or 10 mg/Kg cadmium (Cd) determined using ICP-OES.**

*Columns with different letters are significantly different at 95% confidence interval

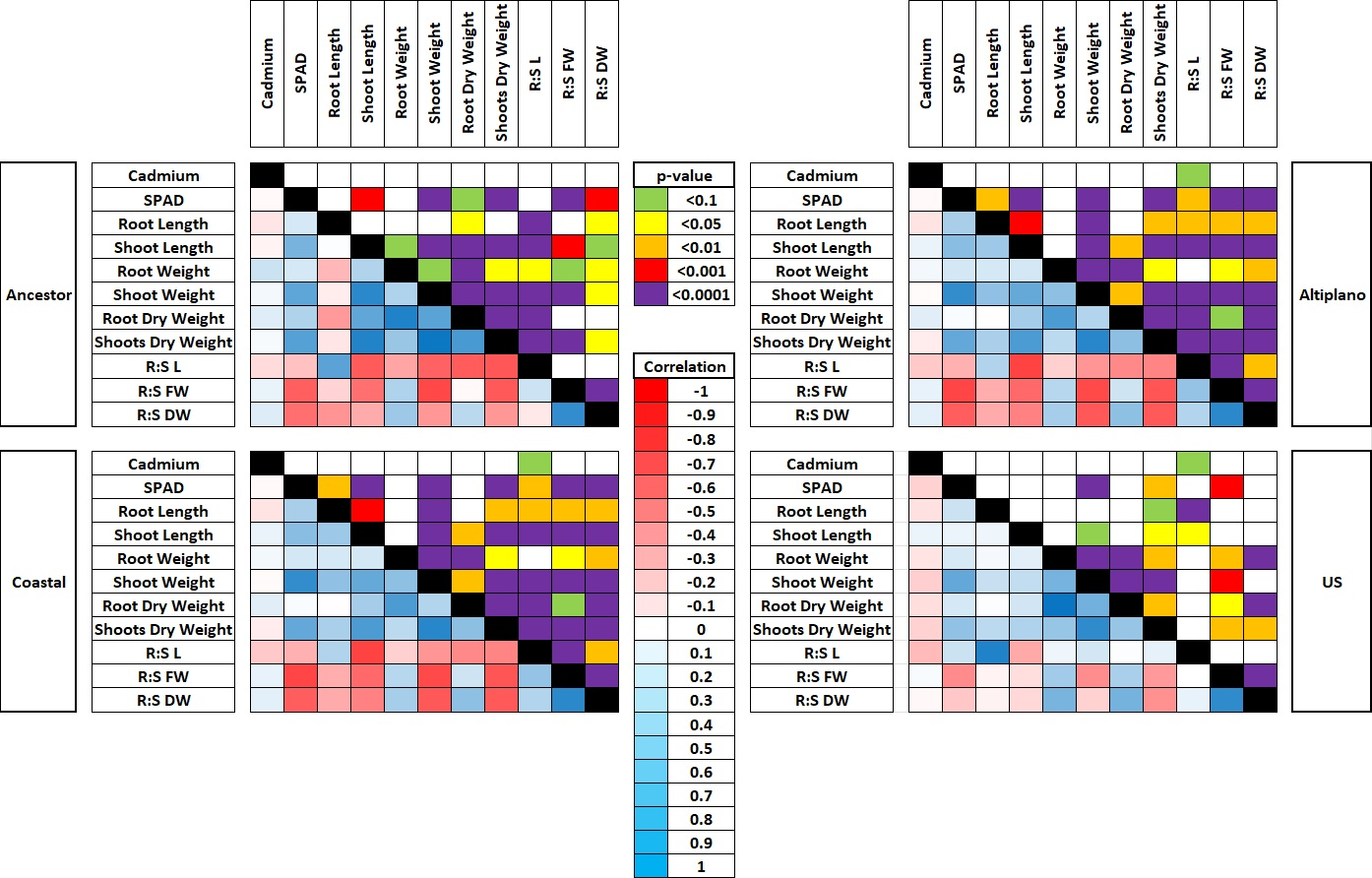

**Figure S1.** **Correlations between cadmium concentrations in the potting media and plant growth parameters at harvest in four Chenopodium ecotypes**

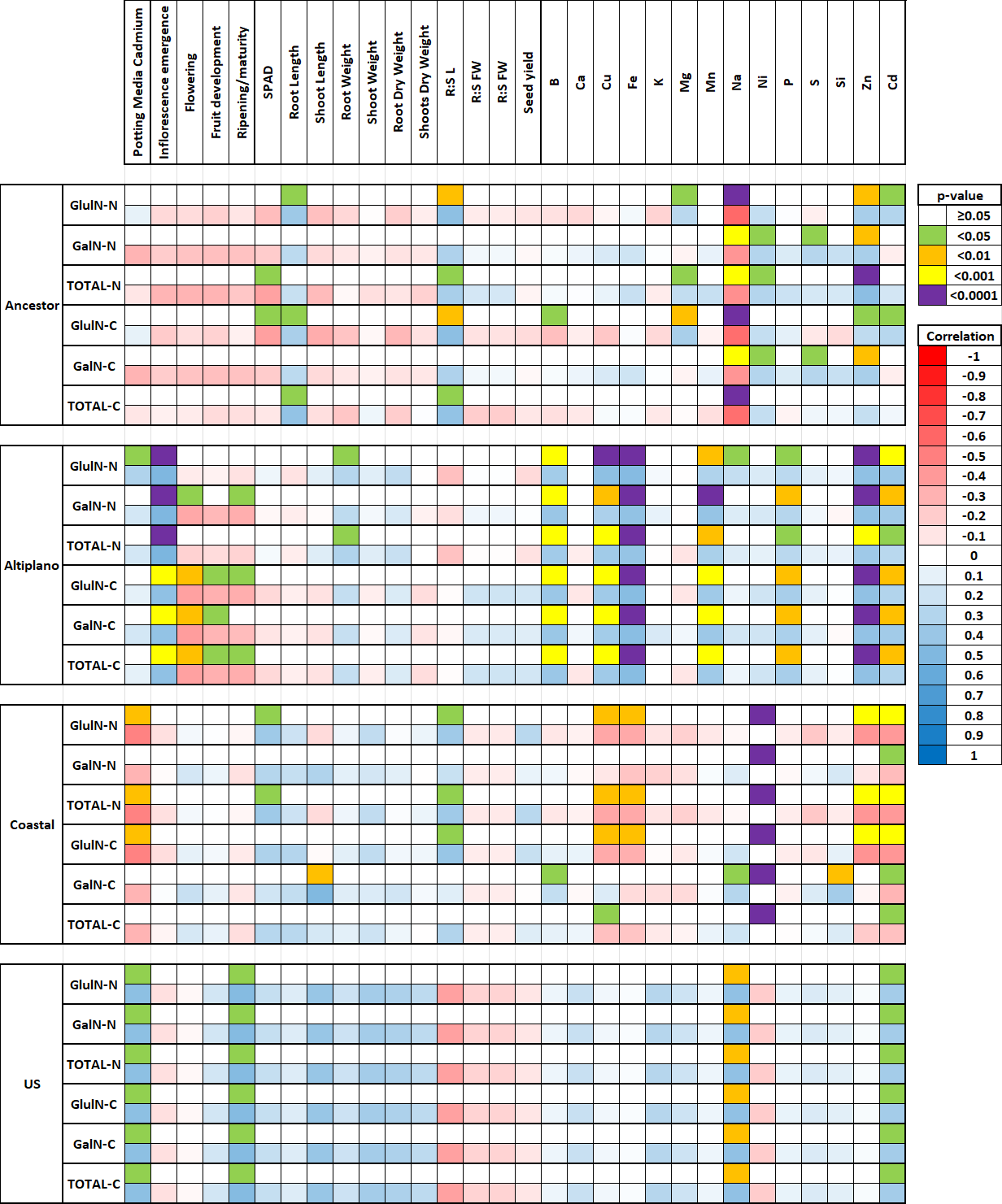

**Figure S2. Correlation of microbial amino sugars with plant growth parameters, plant measurements at harvest, and nutritional profile in the seeds**

Figure S3. a) Isotopic concentration δ^15^N of *quinoa* roots classified by genotype. b) Isotopic concentration δ^13^C of *quinoa* roots classified by genotype.

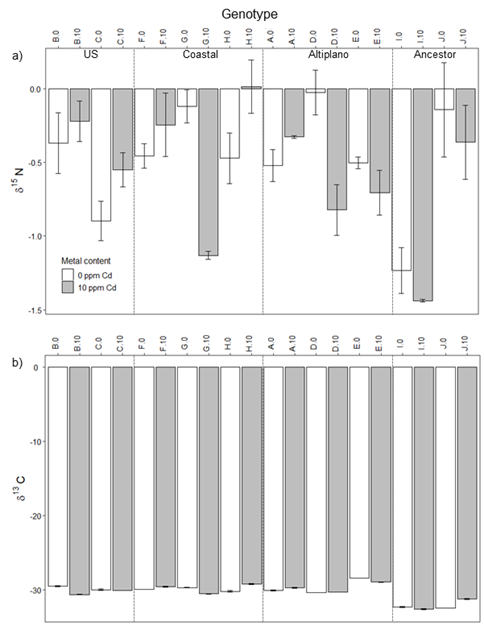
